# Older fathers’ children have lower evolutionary fitness across four centuries and in four populations

**DOI:** 10.1101/042788

**Authors:** Ruben C. Arslan, Kai P. Willführ, Emma Frans, Karin J. H. Verweij, Mikko Myrskylä, Eckart Voland, Catarina Almqvist, Brendan P. Zietsch, Lars Penke

## Abstract

Higher paternal age at offspring conception increases *de novo* genetic mutations (Kong et al., 2012). Based on evolutionary genetic theory we predicted that the offspring of older fathers would be less likely to survive and reproduce, i.e. have lower fitness. In a sibling control study, we find clear support for negative paternal age effects on offspring survival, mating and reproductive success across four large populations with an aggregate N > 1.3 million in main analyses. Compared to a sibling born when the father was 10 years younger, individuals had 4-13% fewer surviving children in the four populations. Three populations were pre-industrial (1670-1850) Western populations and showed a pattern of paternal age effects across the offspring’s lifespan. In 20^th^-century Sweden, we found no negative paternal age effects on child survival or marriage odds. Effects survived tests for competing explanations, including maternal age and parental loss. To the extent that we succeeded in isolating a mutation-driven effect of paternal age, our results can be understood to show that *de novo* mutations reduce offspring fitness across populations and time. We can use this understanding to predict the effect of increasingly delayed reproduction on offspring genetic load, mortality and fertility.

## Media summary

Fathers’ and mothers’ average ages at birth are increasing throughout the developed world, though they are presently still on par with pre-industrial reproductive timing. A child gets most new genetic mutations from its father. New mutations increase linearly with the father’s age. Hence, we can use the father’s age to understand the effect of new mutations on the child. We find that father’s age predicts lower offspring reproductive success in four populations across four centuries: 17-19^th^ century German Krummhörn, Canadian Québec, and Sweden, as well as 20^th^ century Sweden (total sample size > 1.3m).

## Background

The average child carries about 60 genetic *de novo* mutations, which were not present in either of the biological parents’ genomes [1,2]. Of those that are not functionally neutral, most reduce fitness, as random changes to well-calibrated systems usually do [3]. Importantly, *de novo* mutations can reduce fitness more than inherited deleterious variants, on which purifying selection has had more time to act. The older a father is, the more *de novo* mutations his child will carry. This is dictated by the fundamental fact that cell replication engenders errors [4] and male, but not female spermatogonial stem cells replicate frequently, beginning a regular schedule of one division per 16 days in puberty [5].

Kong et al. sequenced the genomes of parent-child triplets and quartets, so that they could pinpoint mutations and their parental origin [1]. They found that a child’s number of *de novo* single nucleotide mutations could be predicted almost perfectly (94% variance explained) by the father’s age at the child’s birth, henceforth *paternal age*. Thus, paternal age appears to be the main systematic driver of varying offspring *de novo* mutation load. Single nucleotide mutations are the most common mutational event, but copy number variants also increase with paternal age; other structural variants tend to come from the father too [6]. Aneuploidies (aberrant chromosome counts) are an exception: they occur more often when older mothers conceive, possibly owing to the prolonged arrest of their ova in the dictyate [2]. Subsequent studies have confirmed the central role of paternal age for mutations [5].

In clinical research, paternal age has proved its usefulness as a placeholder variable for de novo mutations: after initial epidemiological studies reported paternal age effects on autism [7], sibling comparison studies confirmed they were not due to inherited dispositions [8]. Then, exome-sequencing studies corroborated the paternal age effects by directly counting mutations that were not present in either parent’s exome and found a higher mutational burden in autistic children than in unaffected siblings [9]. These findings elucidated disease aetiology both from an evolutionary and a clinical standpoint, by explaining how an early-onset disease linked to very low reproductive success could linger in the face of natural selection.

Given the links enumerated above, paternal age should, via increased mutations, decrease offspring fitness. By fitness, we mean each offspring’s average contribution to the gene pool of successive generations. This contribution can be approximated by the offspring’s early mortality and number of surviving descendants.

However, most paternal age effect studies focus on medical, cognitive and behavioural traits, such as physical and psychiatric disease, intelligence and violent recidivism [8,10-13]. Though many of these traits plausibly affect evolutionary fitness now, it is not always clear how they affected fitness before the 20^th^ century. Moreover, there are scant records on such traits from this time, and old records are not necessarily comparable to modern records. Births and deaths, or baptisms and burials, on the other hand, have been meticulously recorded in churches. Survival and reproductive success were and still are good operationalisations of evolutionary fitness. And fitness is the most ‘downstream’ phenotype of all, in the sense that all non-neutral mutations affect it by definition [14]. This includes mutations linked to well-characterised syndromes such as Apert’s syndrome and autism with early onsets that are often linked to early mortality and non-reproduction [15].

However, few researchers have examined the effect of paternal age on offspring fitness, especially reproductive success. Studies on humans have examined isolated fitness components such as infant survival, longevity, marriage or reproduction in a single population in one place and at one time [16-19]. Some such studies have focused on longevity, which has an ambiguous relationship to evolutionary fitness owing to life history trade-offs [20]. Some have examined the effect of maternal age or birth order, but ignored paternal age [21]. Some focused mainly on environmental explanations, such as decreased parental investment [22], but these are not necessarily sufficient to explain paternal age effects. If they were, the age of the biological parents would not have had negative consequences in a cross-fostering experiment done on wild house sparrows [23]. Owing to variable methodology and sample sizes across studies, we cannot reliably compare findings to find out if they were different because of theoretically meaningful moderators.

## The Present Study

Here we focused on the offspring’s reproductive success, operationalized as number of children who lived at least to an age of 5. To be able to compare all children of a father, we also included the children who did not have any children themselves, even if they died young. Reproductive success is a good predictor of an individual’s contribution to the next generation’s gene pool [24]. But we also separated early mortality, marriage success and number of offspring to examine successive episodes across the lifespan during which natural and sexual selection occur. Based on evolutionary genetic theory, we predicted that in aggregate we would find small, negative effects of paternal age on offspring fitness throughout the lifespan [25]. Some de novo mutations will have very large negative effects, but many more will be (nearly) neutral. In aggregate, on the population level, this implies a small stochastic increase in deleterious effects with paternal age.

However, humans do not time their reproduction randomly. Therefore paternal age effects may be confounded by social and genetic factors [26], associated with both age of reproduction and offspring reproductive success. Because our goal was to isolate *mutation-driven* effects of paternal age, we analysed the paternal age effect within full biological sibships and separated out a between-family effect. This effectively controls for many potential confounds. Full siblings share a parental gene pool, so that genetic load, which accumulated over generations, is distributed across them randomly. Siblings also usually share much of their early environment, and access to resources such as wealth and land. However, social convention may additionally link inheritance to birth order and sex. Therefore, we also adjusted for a number of other social factors that might be linked to increasing paternal age within families, such as birth order and parental loss.

In so doing, we try to accomplish two goals: first, to isolate a potential biological, mutation-driven effect of paternal age on offspring fitness, and second, to compare different populations across different times and in different places, with high statistical power and comparable methodology. Methods

## Populations

To test our hypotheses before the turn of the 20^th^ century, we used genealogies drawn from church records in the Saint-Lawrence valley, Québec (Canada), the Krummhörn (Germany) and four historical Swedish regions. To compare these populations to 20^th^ century Sweden, we used a population-based linkage study from Swedish national health registers.

We used computerized and linked registers of births (and baptisms), deaths (and burials) and marriages to reconstruct family pedigrees and life histories for individuals. We call the individuals whose father’s age we compared with their siblings’ "anchors" wherever it aids comprehension. Further statistics can be found in Table 1 and on the online supplementary website at https://rubenarslan.github.io/paternal_age_fitness/ [27], where all data processing steps and analyses are documented fully.

**Table 1.** Descriptive statistics. RS: reproductive success. IS: infant survival. Numbers in parentheses are standard deviations. Years refer to the birth years of the anchors. For 20^th^-century Sweden, fertility-related numbers are from 1947-1959 (first N given) and mortality numbers are from 1969-2000 (second N given) and the Bayesian reproductive success models were based on a random subset of 100,000 families.

|  | **1720-1850**<br>**Krummhörn** | **1670-1750**<br>**Québec** | **1760-1850**<br>**Sweden** | **20<sup>th</sup>-**<br>**century**<br>**Sweden** |
| --- | --- | --- | --- | --- |
| **Population N** | 80,808 | 459,591 | 271,129 | 8,201,968 |
| **Anchor N** | 16,433 | 107,099 | 187,121 | 1,419,282/<br>3,428,225 |
| **Anchors/ Families**<br>**(RS models)** | 13,989/<br>3,266 | 78,022/<br>12,951 | 84,557/<br>24,595 | 1,207,601/<br>795,812 |
| **Anchors/ Families**<br>**(IS models)** | 16,382/<br>3,762 | 96,042/<br>17,030 | 179,375/<br>49,321 | 3,335,772/<br>1,828,187 |
| **Paternal age** | 35.23 (7.56) | 36.28 (8.48) | 34.37 (7.69) | 31.84 (7.05) |
| **Maternal age** | 31.53 (5.88) | 29.58 (6.66) | 31.54 (6.32) | 28.34 (6.11) |
| **Female infant**<br>**mortality** | 11.1% | 19.0% | 12.0% | 0.5% |
| **Male infant**<br>**mortality** | 12.9% | 23.2% | 14.1% | 0.7% |
| **Fertility (married**<br>**women)** | 3.66 (2.89) | 7.71 (4.57) | 3.6 (3.17) | 2.15 (1.11) |
| **Male age at first**<br>**child** | 29.29 (5.36) | 27.92 (5.29) | 28.13 (5.18) | 28.07 (5.6) |
| **Male age at last**<br>**child** | 39.6 (7.5) | 44.19 (8.59) | 37.52 (8.29) | 33.57 (6.14) |

The first population are the French settlers of the Saint-Lawrence valley in contemporary Québec, Canada [28]. They were an isolated frontier population in a harsh climate but they also had access to abundant resources and lots of unsettled land. We focused on the 107,099 anchors born between 1670 and 1750. Married female anchors from this period had on average 7.7 children.

The second population are inhabitants of the Krummhörn in contemporary Germany [29]. They were also quite isolated and had a stable population size. We focused on the 16,433 anchors born between 1720 and 1850. Married female anchors from this period had on average 3.7 children.

The third population are Swedes in the Sundsvall, Northern inland (Karesuando to Undersåker, includes Sami people), Linköping and Skellefteå regions [30,31]. All individuals living in Skellefteå and most individuals in Sundsvall were linked between church parishes. In the other regions, some individuals appeared in more than one parish. We focused on the 187,121 anchors born between 1760 and 1850. Married female anchors from this period had on average 3.6 children.

Our modern data is the whole population of Sweden. The Swedish Multi-Generation Register includes records of individuals born after 1932 and alive by 1962, as well as their parents. It was linked to the Cause of Death register that includes death dates. Information about marriages was derived from the Longitudinal Integration Database for Health Insurance and Labour Market Studies (LISA by Swedish acronym). Because of data availability and censoring in this dataset, we focused on the 1,419,282 anchors born between 1947 and 1959 for reproductive outcomes and the 3,428,225 anchors born between 1969 and 2000 for survival outcomes. Married female anchors from the earlier period had on average 2.2 children.

## Statistical approach

We employed multilevel regressions with an intercept per family to compare siblings within families. We used the R packages lme4 [32] and MCMCglmm [33], and adjusted for average paternal age within each family to isolate a within-family paternal age effect. In our simple models, which we could run with almost all data, we adjusted only for offspring sex and birth cohort. We adjusted for birth cohort to account for secular changes in mortality and fertility. In the historical Swedish data, we additionally adjusted for geographical region and in the Québec data we adjusted for whether they were born in a major city. In extended models, we added a number of covariates to rule out various alternative explanations. We adjusted for parental deaths in the first 5 years of life to remove effects related to early orphanhood and parental senescence. Because parental death dates were sometimes missing, we had to run these models with slightly reduced sample sizes. We computed the number of living siblings who were younger than 5 years to adjust for a measure indicative of a crowded crib, suggesting dilution of parental care [34]. We adjusted for maternal age in three bins: 20 and younger, 21-34, 35 and older. We binned maternal age to reduce multicollinearity with paternal age, and because it often has nonlinear effects. We also adjusted for family size.

We used MCMCglmm to analyse reproductive success for *all* offspring, including those who died in childhood or never married. By using a zero-alteration link function, we were able to properly model the large number of offspring who died childless. For 20^th^-century Sweden, we had to randomly sample a hundred thousand families to make the MCMCglmm models feasible. We checked our inferences by using different random subsets and by using all data in lme4. Our estimates of the highest posterior densities are thus too conservatively broad for 20^th^-century Sweden. To separate effects into successive episodes of natural and sexual selection, we adjusted for success in the previous episode. For example, to analyse reproductive success we included only ever-married anchors and adjusted for their number of spouses.

## Results and Discussion

We found negative paternal age effects on reproductive success in all four populations, even after adjusting for numerous covariates, including offspring sex, birth cohort, number of (dependent) siblings, loss of either parent, and maternal age. Compared to a sibling born when the father was 10 years younger, anchor individuals had 13% [95% credible interval: 3-22%], 4% [2-6%], 9% [6-12%], and 5% [4-6%] fewer children who survived to age 5 years in the Krummhörn, Québec, historical Sweden and 20^th^-century Sweden, respectively. When assuming an average anchor with average reproductive success for the population, this translates to 0.18, 0.19, 0.20, and 0.10 fewer surviving children per decade of paternal age, respectively (Figure 1).

**Fig. 1:**
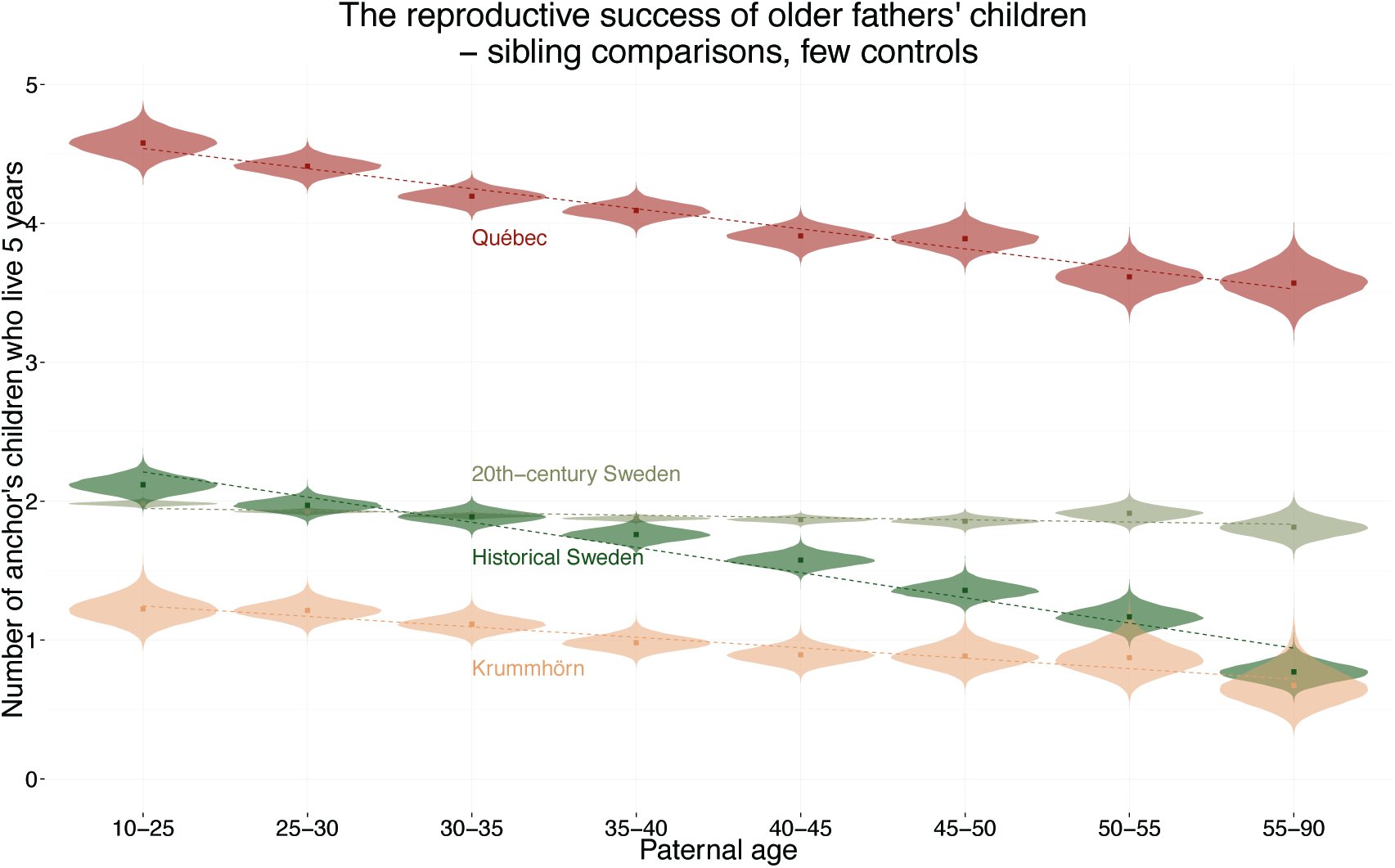
Paternal age effects on number of surviving children. MCMCglmm model predictions for within-family paternal age effects with covariates set to the earliest respective birth cohort and the other covariates, maternal age, parental loss, sex, number of siblings (total and dependent) set to their central tendency. In these heavily controlled models, the following ns remained in the analysis: Québec (51,433), Krummhörn (9,449), historical Sweden (83,850), 20th-century Sweden (1,197,862, plot based on a random subset of 152,032). Points show the posterior mean; the cat eyes show the posterior density (uncertainty). The fitted regression slopes are weighted by certainty.

We observed only effects consistent with a linear dose-response relationship across the paternal age range (Figure 1), as we predicted assuming continuously occurring mutations in the male germ line drove them.

In the Québec data, we had access to deep pedigrees, allowing us to compare not only siblings, but also cousins in a within-extended-family design. Even across three generations, we found negative grandpaternal age effects (maternal grandfather: 4% [2-6%], paternal grandfather: 7% [4-9%] fewer children) on offspring reproductive success, operationalised as above, that were roughly half the size of paternal age effects in the same model (10% [8-12%] fewer children). We find it unlikely that the environment or epigenetics caused these effects, since we would expect such effects to wash out faster than effects based on genetic mutations whose likelihood of being passed on halves every generation.

When we, as early studies have tended to do [17,19], compared individuals *between* families instead of within sibships, paternal age was not consistently negatively associated with fitness outcomes. In fact, we found a positive between-family effect of average paternal age in the models with the basic set of covariates that disappeared when adjusting for further potential social confounds like family size. This suggested to us that our approach succeeded in separating out a non-mutational influence, such as family size or heritable parental fertility dispositions (see supplement [27]).

As shown in Figure 2, when we separately examined the selective episodes along the lifespan, effects were significantly negative across almost all models in Québec and historical Sweden. In the small Krummhörn population (*ns* = 4,433-11,505) only marriage success was significantly negatively predicted by paternal age after adjusting for the extended covariates. Confidence intervals for the other effects still included the estimates from the other populations, but were not consistently significant. This suggests that low statistical power prevented us from breaking up the total effect into the contributions of each selective episode in the Krummhörn.

**Fig. 2:**
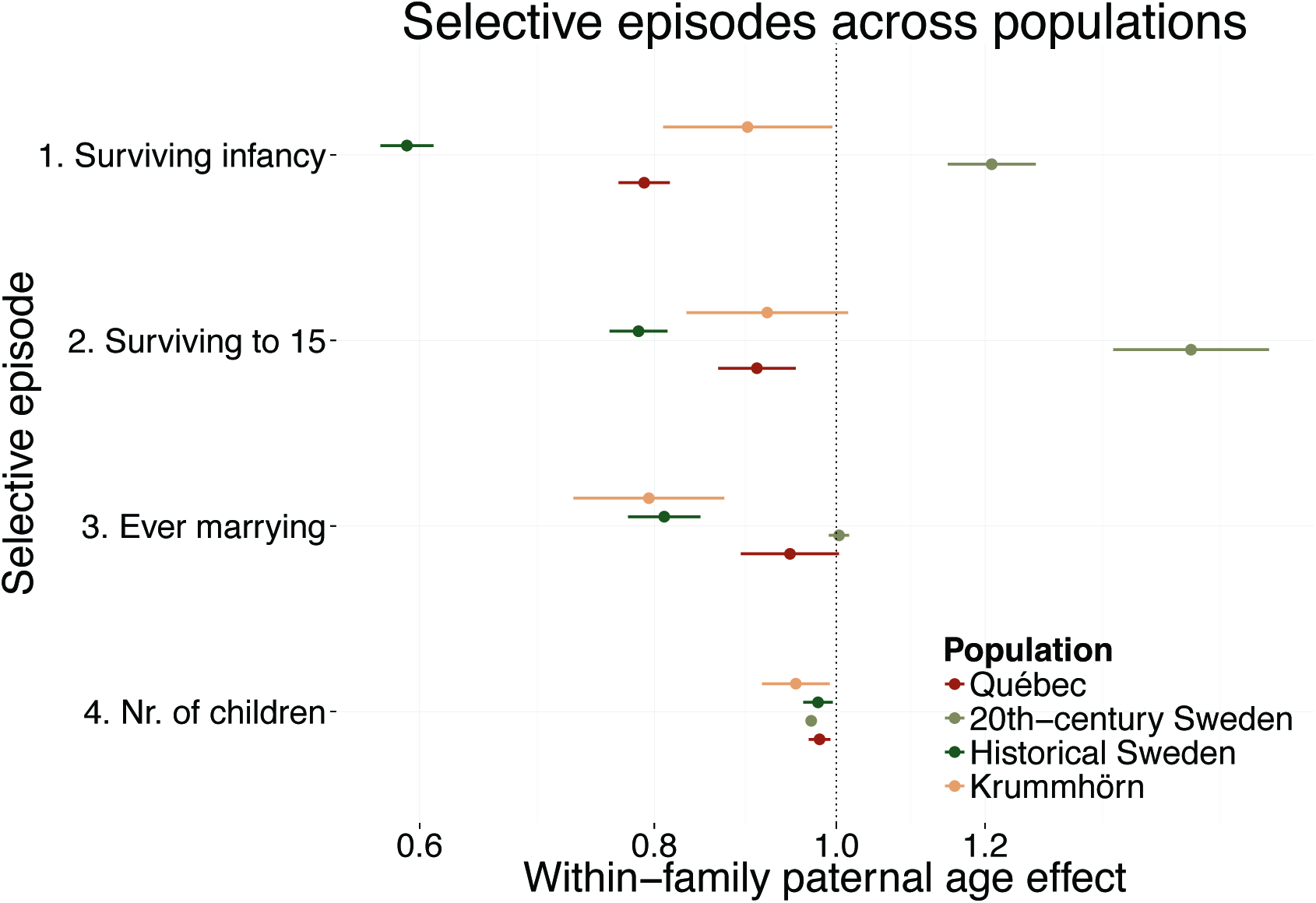
Paternal age effects on subsequent selective episodes. Estimates of odds ratios and intercept rate ratios from lme4 with 95% confidence intervals. Controlled for sex and cohort. Modelling specifics, base rates, *n*s, effects after adding further controls and linearity checks for each of the models are given in the supplement.

In 20^th^-century Sweden, only the effect on number of children was significantly negative. Here, low power cannot explain the absence of negative effects, as this population was our largest by orders of magnitude. Infant and child mortality in Sweden are among the lowest in the world. Because more than 99% of children brought to term in the years 1969 to 1999 survived, natural selection had less leeway to act during this episode. Low infant mortality was achieved mostly through advances in peri-and postnatal care, but abortion may also have played a role. Abortions end one fifth of all known pregnancies in Western Europe [35]. They are unobserved in all our populations. Most are elective, not therapeutic [36], but even women electing to have an abortion may do so selectively after considering their own age and paternal characteristics, including age [37]. On the other hand, paternal age predicts both miscarriages [38] and very preterm births [39]. We can speculate that the selective abortion of fetuses with congenital disorders and the increased survival of preterm births might explain the absence of a negative paternal age effect on infant survival in 20^th^-century Sweden. Unlike surviving to reproductive age, being married may no longer be a prerequisite for high fitness. Although there is still plenty of variability in odds of marriage, you no longer need to marry to start a family in Sweden. For the Swedish generation we examined, born from 1947 to 1959, co-habitation and having children were possible without ever marrying. Unfortunately, we do not have data on romantic co-habitation.

We found it striking how closely aligned the effect sizes were across four populations that were distant in both space and time. This speaks for a universal effect, perhaps inherent in human reproductive biology. With the work of Kong et al. and others [1,5] having demonstrated such a strong and likely causal effect of paternal age on de novo genetic mutations, we are confident to interpret paternal age as a placeholder variable for the occurrence of the latter. However, some alternative explanations for paternal age effects have been put forward and are worth addressing.

Eisenberg et al. [40] linked advanced paternal age to longer offspring telomeres, but it remains unclear whether this association is causal, whether it would differ between siblings and whether it could mediate phenotypic effects. Paternal age effects on epigenetic alterations have so far only been speculated on.

Maternal age is another matter: its effects on aneuploidies are well established in the literature [41]. Although we adjusted for between-family maternal age effects, parents’ ages within families increase in lockstep. Their effects are difficult to separate in largely monogamous populations. Even though maternal age is linked to aneuploidies, most aneuploid conceptions are not carried to term and even live-born children rarely get old. Only children with Down’s syndrome live longer, at least in 20^th^-century conditions, but they are rarely fertile. We observed a linear association between parental age and fitness, consistent with a mutation-driven paternal age effect, but inconsistent with maternal age effects via Down’s, which are typically non-linear, i.e. sigmoidal [41].

Paternal age may itself predict Down’s syndrome when mothers are older [42], but we think this pathway cannot explain all reported effects (see robustness checks). In modern epidemiological data, specific syndromes could be easily excluded to test their contribution. Given that a recent study also estimates a small effect of maternal age on single nucleotide *de novo* mutations [43], we may have been able to isolate a biological effect through our extensive adjustments for social explanations, even though we could not perfectly separate out each parent’s contribution.

In robustness analyses, we tested whether our modelling decisions affected our main results. Our results were robust to a) adjusting for birth order and last born status b) adjusting for age at orphanhood instead of parental loss until age five c) adding separate random intercepts for mother and father instead of one per dyad d) adjusting for birth cohort with a continuous measure or more fine-grained bins e) adjusting for paternal age at first birth in addition to average paternal age in the family f) simulating a larger-than-observed, positive paternal age effect on offspring survival in 20^th^-century Sweden, where some early deaths were not recorded g) adjusting for grandparental loss (where known) in grandpaternal age effect analyses in Québec h) performing survival analyses for the Québec and Krummhörn mortality data and i) randomly omitting childless children of older mothers at several times the actual rate of Down’s syndrome in 20^th^-century Sweden. To find out whether results were driven by first-and last-born adult sons being more or less likely to be the main family heirs, we adjusted for this status. Even though this meant controlling for an intermediate outcome (adult sons necessarily did not succumb to child mortality), results were robust to this adjustment too.

In sensitivity analyses, we showed that effects on fitness outcomes beyond early survival could not be completely explained by offspring education (only available in 20^th^-century Sweden), or offspring reproductive timing (age at first and last birth). However, anchor mortality before 50 years of age seemed to near-completely account for effects on reproductive success in Québec and historical Sweden, but not in the Krummhörn and 20^th^-century Sweden. Age was more often censored than completed fertility, owing to the longer follow-up required. We adjusted for age in four bins (died younger than 25, 25 to 50, died older than 50, death date unknown). The results were robust to binning ages differently.

## Implications and conclusions

We mainly aimed to isolate paternal age as a biological causal factor in our analyses. Indeed, in four large population-based datasets, we find robust support for the evolutionary genetic prediction that higher paternal age linearly decreases offspring fitness via *de novo* mutations. Paternal age effects could also have implications for policy: Descriptive data show a fall from 1930 to 1970 and a steady rise in maternal and paternal ages since 1970 in Sweden. However, average parental ages in 2010 were still lower than in 1737-1880 (Figure 3). Although people start reproducing later, they also stop earlier. Contrary to common news and lay scientific accounts, contemporary parents do not reproduce unprecedentedly late *on average* [1,37]. While advanced parental ages at *first* birth may entail smaller families, pre-industrial populations had similar average ages at birth and were not overwhelmed by mutational stress. So, we do not predict that contemporary reproductive timing will lead to unprecedented or unbearable *de novo* mutational loads. Contrary to oft-repeated doomsaying [44], purifying selection against mutations, in so far as paternal age effects on fitness are an appropriate index, has not been completely cushioned in the age of modern medicine.

**Fig. 3.**
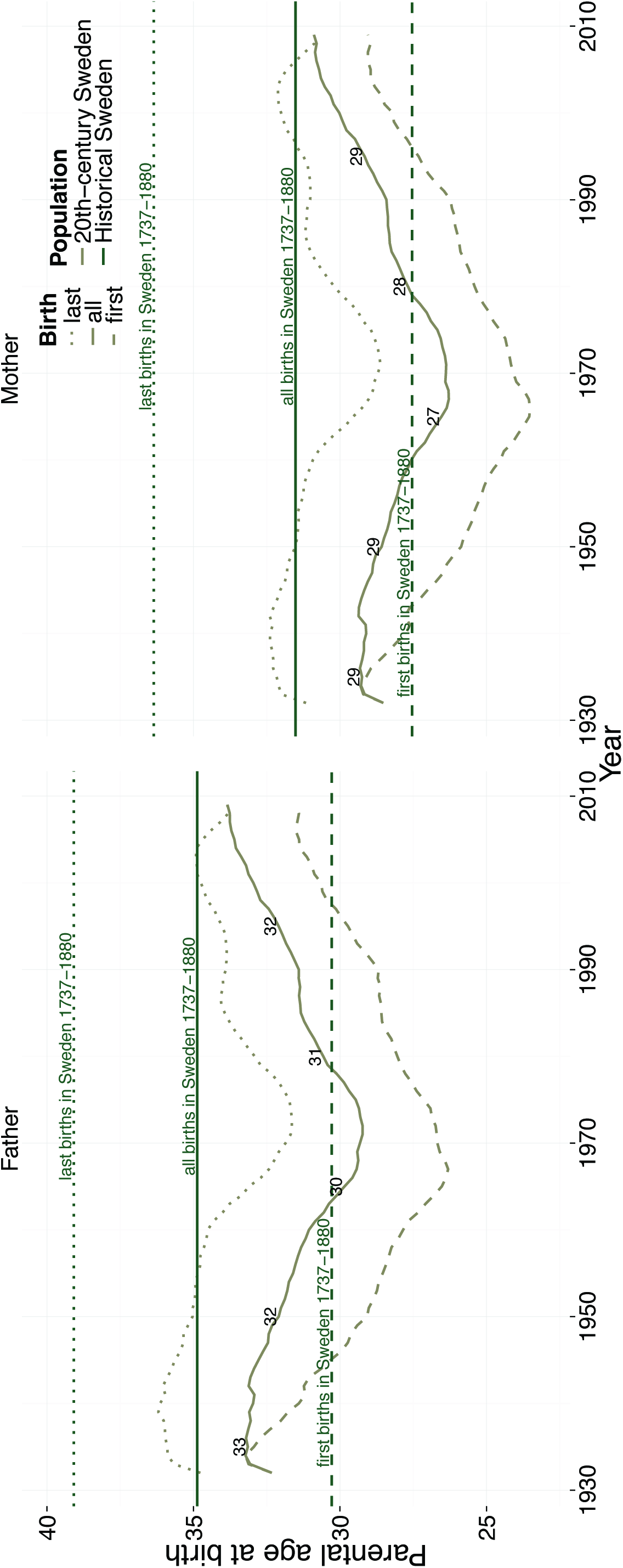
Reproductive timing data showed that average parental ages at birth decreased in 20^th^-century Sweden until ca. 1970 and increased thereafter. Average contemporary parental ages are still lower than in any of the three historical populations. Ages at first birth in the early periods and ages at last birth in the late periods are censored and hence biased towards the age at all births (which is itself unbiased).

Although our design is not ideal for separating the influence of maternal and paternal age, most secular trends and policies will probably affect both. Future research could disentangle the parents’ contributions in polygamous populations or by using genome-sequenced families. Future research could also isolate a biological paternal age effect on early mortality in nonhuman animals with large recorded pedigrees, such as artificially inseminated breeding cattle. This would rule out most social confounds by design, but the much shorter breeding lifespans would limit generalizability to humans

## Competing interests

We have no competing interests.

## Authors’ contributions

RCA and LP conceived of the study. RCA coordinated it, carried out the main data analyses and drafted the manuscript. KPW provided guidance and pre-processing for all church record data, contributed to the design of the study and replicated central analyses in Stata. EF and CA contributed the contemporary Swedish data. EF also provided guidance and pre-processing. EV contributed the Krummhörn data. KJHV, BPZ, MM, and LP helped design the study, helped interpret the data and critically revised the manuscript. All authors helped draft the manuscript and gave final approval for publication.

## Acknowledgements

RCA thanks Jarrod Hadfield, Ben Bolker and Holger Sennhenn-Reulen for their statistical packages and advice. MM acknowledges the support of the European Research Council grant 336475. CA and EF acknowledge financial support from the Swedish Research Council through the Swedish Initiative for Research on Microdata in the Social And Medical Sciences (SIMSAM) framework grant no. 340-2013-5867.

